# Phenomics demonstrates cytokines additive induction of epithelial to mesenchymal transition

**DOI:** 10.1101/2024.05.06.592642

**Authors:** Alphonse Boché, Mathieu Morel, Sabrina Kellouche, Franck Carreiras, Ambroise Lambert

## Abstract

Epithelial to Mesenchymal Transition (EMT) is highly plastic with a program where cells lose adhesion and become more motile. EMT heterogeneity is one of the factors for disease progression and chemoresistance in cancer. Omics characterizations are costly and challenging to use. We developed single cell phenomics with easy to use wide-field fluorescence microscopy. We analyse over 70000 cells and combined 51 features. Our simplistic pipeline allows efficient tracking of EMT plasticity, with a single statistical metric. We discriminate four high EMT plasticity cancer cell lines along the EMT spectrum. We test two cytokines, inducing EMT in all cell lines, alone or in combination. The single cell EMT metrics demonstrate the additive effect of cytokines combination on EMT independently of cell line EMT spectrum. Single cell phenomics is uniquely suited to characterize the cellular heterogeneity in response to complex microenvironment, and show potential for drug testing assays.

## 1. Introduction

The Epithelial to Mesenchymal Transition (EMT) is a reversible physio-pathological phenomenon often found in ovarian cancer processes (late stage diagnosis with survival rate reduced to 31.5%) where cohesive tissue with well-polarized cells switch to less cohesive, more motile cell behaviours.^[1,2,3,4]^ This transition is on a spectrum^[5]^ and plays a role in the cellular heterogeneity and therapy resistance in ovarian cancer.^[6]^ EMT can be induced *in vivo* in the microenvironment by particular cytokines or extracellular matrix macromolecules like Fibronectin.^[7]^ Occurring in 40% of advanced stage ovarian cancer, ascites is a type of complex microenvironment occurring by an accumulation of peritoneal fluid which can sometimes be associated to reduced survival and increased dissemination.^[8]^ The presence of cytokines mixture with synergy, antagonist or additive effect on EMT in ascites may explain these outcomes.^[9–12]^ However, characterizing molecules combined effect on EMT remains a challenge.

EMT can be tracked with transcriptomic methods, which are costly and don’t integrate all parameters, including the morphological aspect or single-cell variation.^[13]^ Each cell, even in homogenous condition, can be modified or modify itself from its population.^[14]^ Single cell variation is a necessity in organism, but also a major issue when it comes to treatment, especially in highly heterogeneous malignancy like ovarian cancer. EMT is a heterogeneous and plastic process for which the amount and precision of data needed can be demanding in time or resources.

To address these challenges, we select four ovarian cancer cell lines covering the EMT spectrum.^[15]^ We use phenomics to analyse both morphological and molecular modifications.^[16,17]^ Furthermore, we implement single-cell analysis to describe the intra- and inter-cellular heterogeneity of the EMT.^[18]^ We analyse over 70000 cells among the four ovarian cancer cell lines. An image based analysis entails the single-cell morphology which encodes the metastatic potential^[19]^ and only two single cell EMT molecular markers.^[20,21]^ We designed an algorithm to define unsupervised clustering of phenotypic parameters extracted. Clustering discriminates the four cell lines as a function of EMT spectrum. and discriminates the cytokines responses within each cell line. Based on single cell clustering distance, we define a single parameter of induction of EMT. This simple metric of EMT induction analysis demonstrates that TNFα and TGFβ1 play an additive role in the EMT plasticity in our conditions. We show that with a simplistic pipeline, we can analyse with the highest precision and relatively low entry cost, one the most heterogenous process tuning the cell response in cancer.

## 2. Results

### 2.1 Single-cell image-based simplistic pipeline discriminate EMT states

We based our analysis on four ovarian cancer cell lines spread across EMT spectrum from more epithelial to mesenchymal: OVCAR3 cell line is an epithelial one, IGROV1 cell line a hybrid-epithelial (an epithelial cell line presenting mesenchymal characteristics), SKOV3 cell line a hybrid-mesenchymal (a mesenchymal cell line presenting epithelial characteristics), and A2780_C10 cell line mesenchymal.^[15,22]^ To induce EMT, we used two different cytokines: TNFα and TGFβ1. Qualitative observation of images (**Figure 1A** and **Figure S1**) indicates that TGFβ1 induces the strongest mesenchymal phenotype. After TGFβ1 induction the cells increased their isolations in all cell lines, except for OVCAR3 cell line. OVCAR3 cells had an increase in cortical actin after TGFβ1 induction (Figure 1A and S1). The elongation of cell is also visible particularly in IGROV1 and SKOV3 cell lines in TGFβ1 condition. On the contrary, TNFα seems to induce a Mesenchymal to Epithelial transition, as we observed an improvement in cell-cell junctions.

**Figure 1:**
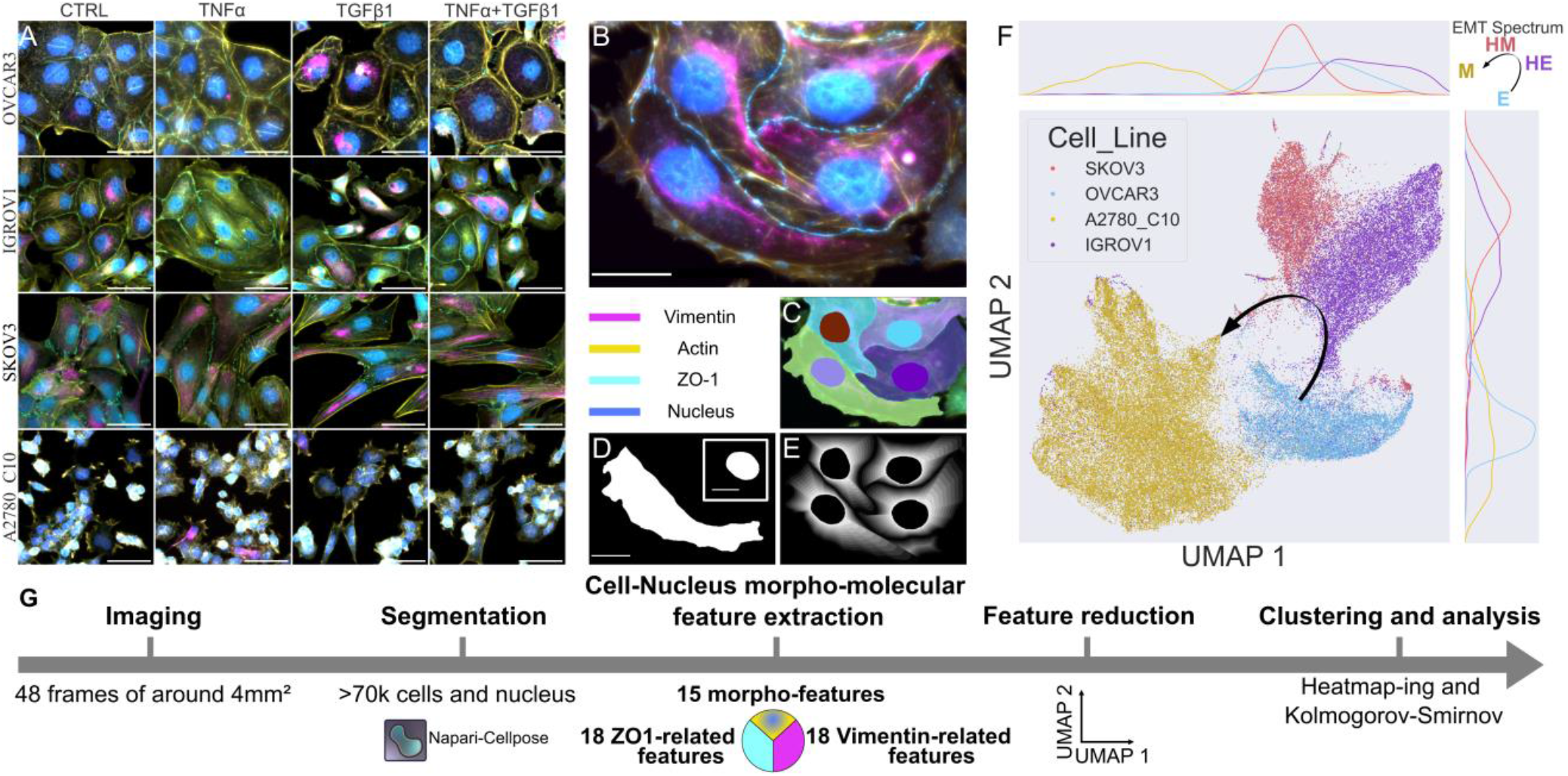
Epithelial to Mesenchymal Transition (EMT) Simplistic Workflow enable efficient clustering of EMT spectrum of ovarian cancer. (A) Four ovarian cancer cell lines: OVCAR3 (Epithelial cell line: E), IGROV1 (Hybrid-Epithelial cell line: HE), SKOV3 (Hybrid-Mesenchymal cell line: HM) and A2780_C10 (Mesenchymal cell line: M). The induction of EMT is induced by the cytokines TGFβ1 and/or TNFα. (B) Along with cellular and nuclear morphologies, EMT parameters include Vimentin (Mesenchymal marker) and ZO-1 tight junction protein (Epithelial marker). (C) Each cell and nucleus are labelled with cellpose (D) leading to a PCA-based method analysis with Celltool on the cell/nucleus morphologies and (E) the normalization from nucleus to cellular membrane. (F) UMAP clustering of ovarian cancer EMT single cell analysis. (G) Pipeline summary: N=3 replicates, 70835 cells in total have been analysed on 51 morphological and molecular features. Scale bar = 50μm (20μm for nucleus in (D)).

We aimed to measure quantitatively the EMT in these different cell lines on an image based and single cell approach. Over a ten thousand cells have been analysed for each cell line, and over thirty thousand for A2780_C10. We labelled vimentin as a mesenchymal marker, an intermediate filament involved in the remodelling of cell shape during EMT. We labelled ZO-1 as an epithelial marker, a tight junction protein (Figure 1B). Finally, we labelled actin and visualized the nuclear membrane through nuclear DNA staining to make a single cell morphology Principal Component Analysis (PCA) on cellpose (Figure 1C).^[23]^

A macro on FIJI^[24]^ has been created to merge the different measurement processes in a semi-automatic approach where the whole analysis is made by itself (script in supplementary). The labels of actin and nucleus were merged and ordered from each cell to each nucleus in a single Region Of Interest (ROI) Manager. The analysis can be started once each cell contours have been automatically checked and manually corrected when needed.

Segmentations generated by cellpose were binarized on FIJI and analysed with celltool.^[25]^ Celltool created a mean morphology, around which we obtained the distribution of morphological deformations within our cell populations for each cell line (Figure 1D). We linked any nucleus data to its cell data, since Nucleus to Cell Ratio is an important factor for *in vivo* malignancy.^[26]^ In addition, we created a normalized topography from each nucleus to its cellular membrane to extract internal molecular distribution (Figure 1E). To do so, we started on the idea to connect with the shortest distance two separated ROIs on FIJI.^[27]^ Then, we modified the original code to do the same, but with one ROI (the nucleus) being within the other one. Afterward, the algorithm will draw the shortest distance with the nucleus for each cellular contour pixel and interpolate each draw to create the different internal ROI for each cell (see detailed script in supplementary). This pipeline led to a characterization of 51 morpho-molecular features extraction for each cell (Figure 1G). We used UMAP for the dimension reduction.^[28]^ We created a clustering of our cell lines in a counterclockwise EMT spectrum along the UMAP parameters (Figure 1F).

### 2.2 Phenotypic parameters analysis after EMT induction encodes EMT signature in ovarian cancer

To compare the cellular and nuclear morphologies, we did a celltool shape analysis for each cell line and each condition at a time (**Figure 2A**). We obtained the average morphology of actin and nucleus contours per cell line and per condition. We described the standard deviation above or below the average cell, along with the percentage of variation it describes for the size and the elongation, as they are the two main deformations observed. Celltool shape analysis highlighted a tendency of ovarian cancer cell lines to increase in size in presence of TGFβ1. Furthermore, when the cell line has a more epithelial spectrum the TGFβ1 increased the morphological deformation (except for the cellular elongation of OVCAR3 cell line). The mean morphology of TGFβ1 and TNFα cytokines combination seemed to correspond to the addition of TGFβ1 and TNFα condition mean morphologies. We represented every parameter used for single-cell phenomics analysis in a heatmap (Figure 2B). Details of the parameter are given in **Table S1**. We made the difference between the mean of the control to every condition, and normalized the result between condition, then between replicates. Parameters included 13 morphological features and for a molecular aspect, 18 features for Vimentin and 18 features for ZO-1. Each cell line had a specific heatmap signature (Figure 2B). Morphological features analysis indicated similar values in TGFβ1 conditions on average over all the features (heatmap value above 0.5) with few effects of TNF-alpha (heatmap value below 0.5), except for IGROV1 cell line. In IGROV1 cell line, TNFα or TGFβ1 had respectively an effect on the morphology (heatmap value above 0.5).

**Figure 2:**
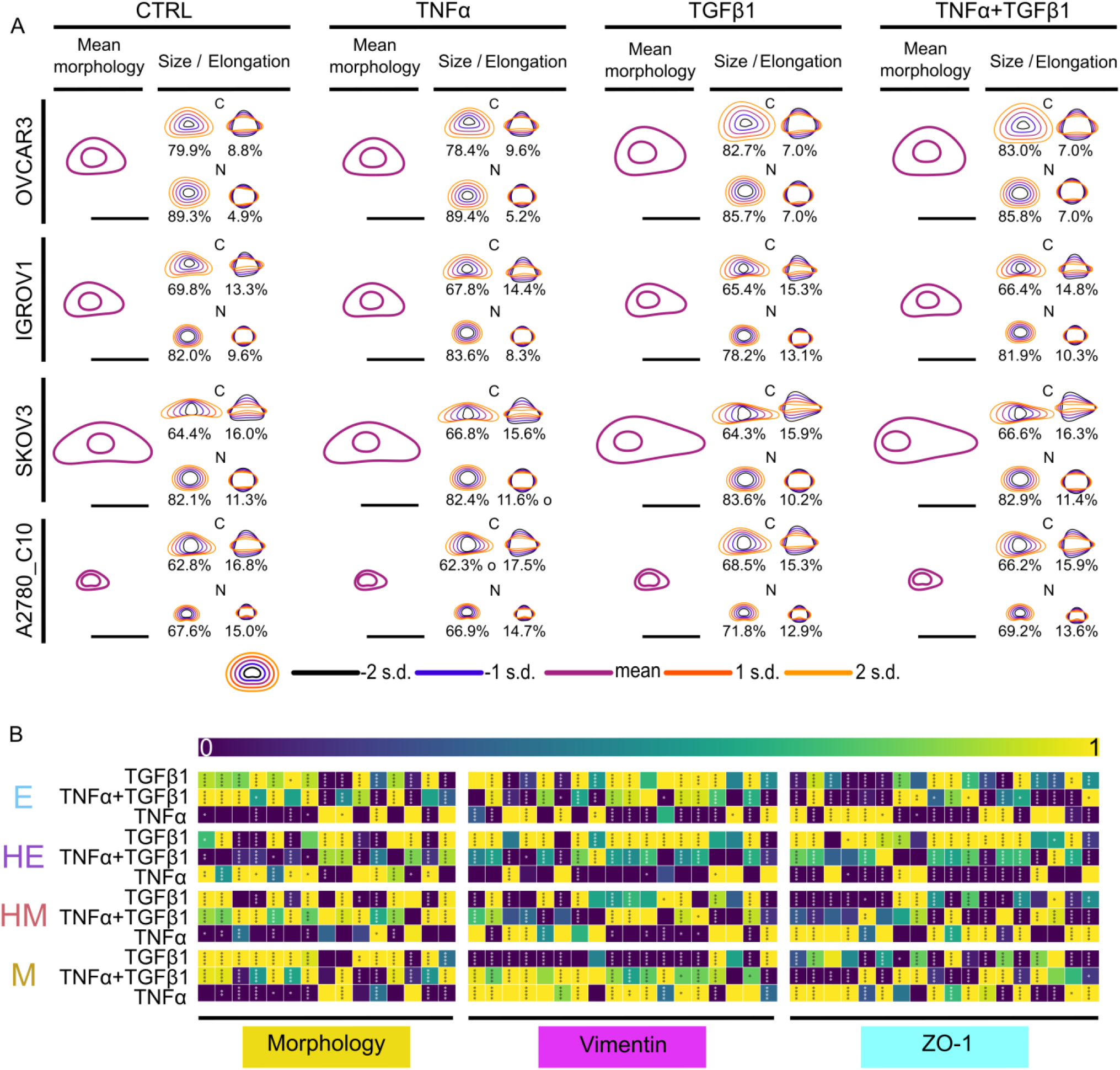
Phenomic defines EMT signature under different conditions for each cell line. (A) Cellular and Nuclear mean morphologies for every condition. The standard deviation of deformation around the mean shapes are represented for both size and elongation (yellow indicates bigger and/or more elongated shapes, black represents smaller and/or rounder shapes). (B) Every parameters used in the UMAP. Each square represents the difference between the mean of the condition and control. Within a cell line, for each parameter (see details in Table S1), the heatmap corresponds to the normalized mean of each replicate normalized mean, among conditions. The significance is related from condition to the control. N=3. *p<0.05; **p<0.01; ***p<0.005;****p<0.001. Scale bar =

For the molecular features, Vimentin (Figure 2B, mid panel) and ZO-1 (Figure 2B, lower panel), OVCAR3 cell line reacted more to TGFβ1 for Vimentin, and more to TNFα for ZO-1. IGROV1 cell line is TGFβ1 driven for both vimentin and ZO-1. SKOV3 cell line is TNFα driven except for the nucleus part of Vimentin where TGFβ1 is more impactful. A2780_C10 cell line is TNFα driven. We can conclude that TGFβ1 alone induces higher modification in epithelial cell lines OVCAR3 and mostly IGROV1, while TNFα alone induced higher modification in mesenchymal cell lines SKOV3 and mostly A2780_C10. Cytokines combination induced an intermediate phenotype between TNFα and TGFβ1 in each group of features (Figure 2B). Furthermore, in all cell lines but especially in IGROV1, the cytokines combination parameters values are included between TNFα and TGFβ1 more than half of the time.

### 2.3 Single-cell phenomics demonstrate additivity of cytokines combination

We then investigated the single cell response to combination of TNFα and TGFβ1. UMAP clustering allows single-cell comparison.^[28]^ UMAP uses dimensionality reduction to understand multiparametric single cells variations (**Figure 3A**). In all the cell lines (OVCAR3, IGROV1, SKOV3, A2780_C10), the distribution of UMAP parameter 1 and 2 depends mostly on the presence of TGFβ1. In epithelial cell lineage, TGFβ1 effect induces more differentiation of single cell phenomes based on UMAP parameters. In OVCAR3 and IGROV1 cell lines, we have a differentiation of the distributions with and without TGFβ1 for both UMAP Parameters.

**Figure 3:**
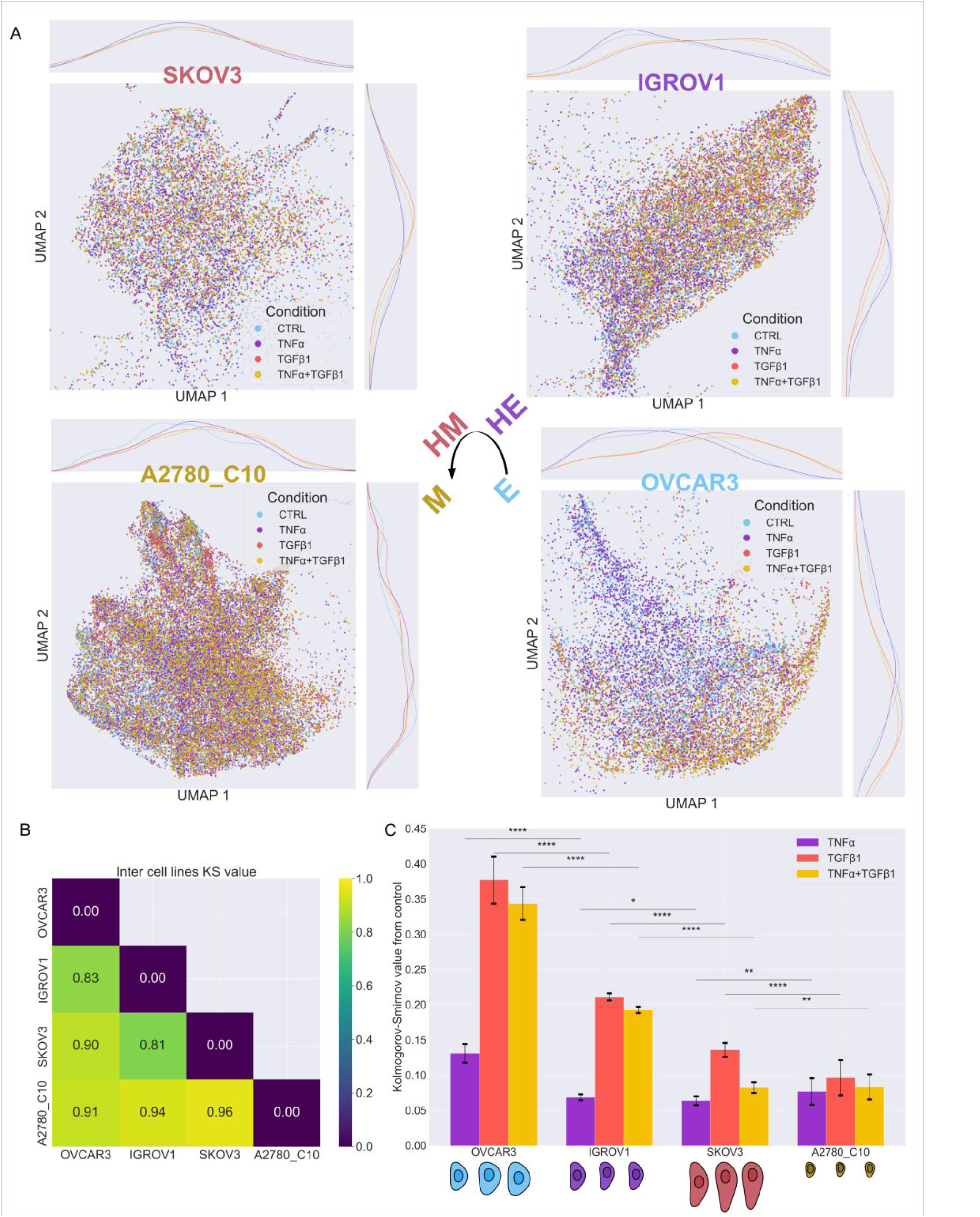
Additivity effect of TNFα and TGFβ1 for the EMT in ovarian cancer. (A) Representation of the UMAP for the four conditions within each of the four ovarian cancer cell lines (Control: cyan / TNFα: purple / TGFβ1: orange / TGFβ1 and TNFα: yellow). The kernel distribution of the conditions for each UMAP Parameter are also represented. (B) Kolmogorov-Smirnov (KS) values between the four ovarian cancer cell lines. (C) KS values between each cytokines conditions and control for each cell line. The mean morphology for each condition is represented below its histogram. N=3. KS values are obtained from the mean of 50 UMAP generation for each replicate. The error bars are the mean of the standard deviation of each replicate. *p < 0.05; **p < 0.01; ***p < 0.005; ****p < 0.001

We observe it in SKOV3 cell line only for the second UMAP parameter. UMAP Parameters distributions are highly heterogeneous in A2780_C10 cell line, but we observe a more distinct separation of the TNFα from the control compared to the other cell lines.

We can merge the UMAP clustering with any single parameter. We create a single parameter heatmap fitted on the UMAP clustering of each cell line (**Figure S2**). If a parameter is relevant in the cellular clustering, the heatmap will be more homogeneous along UMAP Parameters. By comparing the different group of features (morphology, Vimentin and ZO-1), we can determine which aspects are relevant in the EMT induction of each cell line. UMAP parameters are cell line dependant for each pipeline parameter. As expected from the UMAP distributions presented in Figure 3A, OVCAR3 and IGROV1 cell lines present more homogeneous heatmaps for the different features than SKOV3 and A2780_C10 cell lines (Figure S2). OVCAR3 cell line has continuous heatmap for all three group of features. IGROV1 cell line is more homogeneous in the distribution for Vimentin and ZO-1 features than on the morphological features. SKOV3 cell line presents a more inhomogeneous heatmap on molecular aspect and a more homogeneous heatmap on morphological aspect. A2780_C10 cell line has homogeneous heatmap for both morphological and molecular feature. Our results show qualitatively the relevance of each parameter in the UMAP clustering. By comparing the heatmaps of parameters with the distribution of conditions in Figure 3A, we can obtain the most relevant parameters to the EMT induction for each cell line. Finally, we can display a dynamic cell atlas view (supplementary movie 1).

For each cell line in Figure 3A, the distribution of conditions is represented. Most of the time, if the TNFα distribution is above the control distribution, the TNFα and TGFβ1 combined condition is above the TGFβ1 condition. Same goes if the distribution is below. To compare the different inter-conditions clustering, we use the Kolmogorov-Smirnov test.^[29]^ This test establishes an empirical difference between two clusters (**Figure S3A**), with a KS value of 0 for two undistinguishable clusters and a KS value of 1 for two perfectly separated clusters. UMAP are generated randomly, leading to different distances between model iteration. The KS values between cell lines are all above 0.81 and up to 0.96 from over 50 UMAP generation for each of our replicate (Figure 3B). It indicates a robust distinction between our cell lines by the UMAP clustering. Similarly, for each cell line, we compared the KS value from our cytokines conditions with the control from over 50 UMAP generation for each of our replicate (Figure 3C). The KS value, in our case, represents the effects of conditions on single cell EMT plasticity. Three cases are possible when comparing the KS value of the combination of TNFα and TGFβ1 to the single cytokine conditions: antagonistic, additive or synergistic.^[30]^

The combination of TNFα and TGFβ1 would have an antagonistic action if the effect is lower than any cytokine alone:

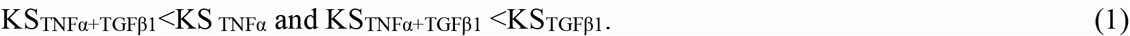

The action would be additive if the effect is intermediate between TNFα and TGFβ1:

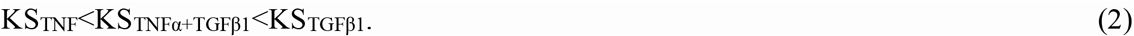

Finally, the action would be synergistic if the effect is higher than any cytokine alone:

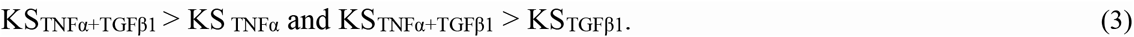

Our results demonstrated that for epithelial cell lines, cytokines affected more the EMT program. Our results also highlighted that the KSTNFα+TGFβ1 is for each cell line between the single cytokine conditions (**Equation 2**). We compared the KS value from TGFβ1 to the three other conditions (Figure S3B). The KS value obtained for all cell lines indicate a constant of around 0.06 for TNFα and TGFβ1 condition. In this case, TNFα condition had an equivalent or higher KS value than control from TGFβ1 condition. We can conclude that the combination of TNFα and TGFβ1 is additive for the EMT induction in ovarian cancer.

## 3. Discussion and Future Perspectives

Image acquisition and analysis is costly in efforts and time, which are precious resources.^[31]^ We design an efficient, free and easy to use algorithm allowing to generate an image based phenomics analysis right after the cellular segmentation process. Our pipeline is developed on FIJI as it is broadly used in life science research. It assures a semi-automatic analysis for unaccustomed researchers. It is a modular resource, that can be integrated to complete different analysis routine. We indexed cell as a function of their nucleus to achieve a single-cell-and-nucleus image-based analysis. To our knowledge, we included one of the first intra-cellular topology tool: we discretize the cytosolic area as a function of nucleus to plasma membrane distance. It allows the localisation of internal features from nucleus to cell membrane, which is one of the most discriminating parameters for our Vimentin data (Figure 2B). Our parameters combined distinguish with precision our four cell lines.

Cytokines are cellular signalling molecules and have a role in the cellular life and fate.^[32]^ Cytokines act on multiple signalling pathways. TNFα and TGFβ1 cytokines can be pro or anti-inflammatory depending on the cell state and its microenvironment.^[33]^ These cytokines have been reported mostly in synergistic and antagonistic effects.^[34,35]^ TNFα and TGFβ1 combination induce an antagonist reaction for tissue regeneration and wound healing.^[36,37]^ Tissue repairing is also part of the EMT process with TGFβ1.^[38]^ TGF-β2, another member of the TGF family, has shown antagonist effect with TNFα treatment ^[39]^. Several studies pointed out the synergistic effect of TNFα and TGFβ1 on EMT, in colonic organoids or breasts cancer.^[10,40]^ It is therefore important to mention that on our exposure time, we found no study describing the combination of TNFα and TGFβ1 in ovarian cancer. To the best of our knowledge, we report for the first time additive effects on EMT of TNFα and TGFβ1 in any cancer type.

EMT is known as a biological process involving molecular and morphological modifications.^[4,5,41]^ Individually taken, our parameters or features could express synergy or antagonism. Here, molecular features variability (Vimentin and ZO-1) explains better IGROV1 cell line plasticity and morphological features variability (cell and nucleus) explain better OVCAR3 cell line plasticity. Our image based phenomics pipeline aimed to merge these two types of features at once, on a single cell approach. Indeed, we highlight the additive effect of TNFα and TGFβ1 on EMT in ovarian cancer. Interestingly, based on the clustering distributions, TNFα acting as a pro-epithelial cytokine in ovarian cancer could be further explored.

In complex microenvironment like ovarian cancer ascites multiple cytokines are often found.^[42]^ Their combination and effect are less clearly understood and remain challenging due to multiplicity of parameters and chronicity of the microenvironment.^[8]^ Our approach consisted of checking whether TNFα and TGFβ1 acted individually and/or synergistically to the EMT of different ovarian cancer cell lines. This study is a referential on a minimal parameters EMT spectrum phenomics analysis. This phenomics approach could extend to include the impact of the microenvironment as well. Complex microenvironment like ascites in ovarian cancer can modulate the cell response.^[43]^ We can note that cytokines combination additive effect was demonstrated based on clustering of over 50 single cell parameters. All parameters were represented with a single parameter representing EMT induction (the KS value). This methodology could be tested for different drug combination and drug testing strategies. Cell culture in ascites condition could mimic *in vivo* microenvironment in malignant ovarian cancer cell development.^[44]^

We presented a semi-automatic workflow for an image-based analysis. It ensures that based on a segmented microscopic image, the analysis would be done under relatively low supervision. The manual correction is essential to generate qualitative segmentation. It constitutes the ground truth estimation, that would enable Convolutional Neural Network (CNN) implementation. CNN could be implemented to obtain a fully automated pipeline from the microscopic image to the data analysis. Different solutions exist already, like training a CNN without annotation or with existent ones.^[45,46]^ In this case the nucleus segmentation, which is easily created by current automatic segmentations tools. Integrating automatic segmentation to our pipeline would increase the efficiency of the workflow.

Dynamic approaches of static snapshots during a period, could also be relevant. Phenomics is a promising multi-omics approach for scientific and clinical field.^[16,47]^ Existent work focus on the integration of phenomics imaging to predict clinical outcome.^[48]^ A similar project exist for the description of EMT in lung cancer through a single cell mass cytometry.^[49]^ Additional imaging modalities, like infrared microscopy allow to correlate single cell information with metabolic state.^[50]^ We believe our pipeline would enable ease of use integration of metabolic state with EMT state. Lately correlation have been established between Warburg effect and EMT.^[51]^ Our pipeline will be a new step to determine the best treatment based on the tumoral clustering of cells from a patient through imaging in vitro.

## 4. Take Home Message

The key factors in this article are first, the creation of a time and cost-efficient semi-automated algorithm for microscopic imagery. By encompassing different molecular and morphological features, we realize a cluster of the EMT program in four representative ovarian cancer cell lines. Second, the additivity of two cytokines, TNFα and TGFβ1, was shown along all four of the studied cell lines. By improving the cell segmentation process, we prospect for this analysis pipeline to be suited for large-scale screening based only on imagery.

## 5. Experimental section

### Cell Lines

IGROV1 is a cell line from a patient diagnosed with a grade III ovarian cancer and has been generously given by the Dr. J. Bénard.^[52]^ The OVCAR3 (ATCC, HTB-161™) cell line is isolated from ascites from an ovarian cancer patient, the A2780_C10 cell line has been established from tumor tissues from an untreated patient and both have been generously given by the Dr. L. Poulain (Unité de Biologie et Thérapies Innovantes des Cancers Localement Agressifs (BioTICLA), Caen). SKOV3 cells derived from malignant ascites of serous ovarian adenocarcinoma were from American Type Culture Collection, Manassas, USA (ATCC, HTB-79 ™).

### Cell Culture

The cells were cultured at 37°C in a humid atmosphere containing 5% CO2. The culture medium is changed every 2 or 3 days, and the cells are passaged twice a week. The cells are detached from the surface by treatment with Trypsin-EDTA 0.25% (Gibco, Ref-25200056) for 5 minutes at 37°C. All experiments were performed using sub confluent cells cultured in flasks of 75cm^2^. The cell culture medium is RPMI-1640 GlutaMAX (Gibco, Ref-72400021) containing 10% fetal bovine serum (FBS, Thermo Fisher Scientific), 1% of penicillin (200 μg/mL) and streptomycin (200 μg/mL) antibiotics (Thermo Fisher Scientific), and 0,075% sodium bicarbonate (Thermo Fisher Scientific).

### Single cell acquisition

#### Cell culture experiment

During experiments, for each cell line, four P4 (Nunc™ Cell-Culture Treated Multidishes, 176740, Thermo Scientific) were filled with a seeding density 10500 cells/cm^2^ for IGROV1 and for SKOV3, 16000 cells/cm^2^ for A2780_C10 and 21000 cells/cm^2^ for OVCAR3 over a 13mm diameter coverslip. These densities are made so cells don’t reach confluence before the end of the experiment to help the segmentation process. After seeding, cells are manually and cautiously shacked on a flat surface without vibrations to prevent cells to aggregate in early process and make a homogeneous distribution of cells within the well.

Cells are at fist cultured in 500μL of medium with 10% (v/v) FBS for 24 hours. The next day, medium is carefully removed and replaced with 500μL of medium with 0% (v/v) FBS. Each P4 medium will contain, except one as control, a concentration of 10ng/mL of TNFα, a concentration of 10ng/mL of TGFβ1 (R&D Systems,240-B-002/CF), or TNFα (R&D Systems, 210-TA-005/CF) and TGFβ1 at 10ng/mL each. The carrier free cytokines were both resuspended following the R&D Systems recommendation. After 24 hours, each medium is renewed with a similar condition for another 24 hours before immunolabeling.

#### Immunolabeling

The cells on coverslips were first washed three times with Phosphate-Buffered Saline (PBS, Sigma-Aldrich, P4417) before being fixed with a 4% (v/v) Paraformaldehyde treatment (PFA, Sigma-Aldrich) for 10 minutes. After fixation, the cells received 3 washes of PBS before being treated with Triton-X (0.1% (v/v) in PBS) at 4°C for 5 minutes. This was followed by 2 washes with Bovin Serum Albumin (BSA, Thermo-Fisher, A7906) at 0.5% (v/v) in PBS and then placed in BSA at 1% (v/v) in PBS for 45 minutes for saturation. After this saturation step, the coverslips were placed on parafilm positioned on the bench, cells exposed. The primary antibody solution was prepared with a dilution of 1:100 of both rabbit anti-human anti-ZO-1 (Sigma-Aldrich, AB2272) and mouse anti-human anti-Vimentin (Sigma-Aldrich, CBL202) in 0.5% (v/v) PBS-BSA, and 20μL was placed on each coverslip for 1 hour under a humid dark chamber. Each coverslip is then put back to its original well to receive three washes with 0.5% (v/v) PBS-BSA. The secondary solution was prepared with a dilution of 1/100 of goat anti-mouse Alexa Fluor 700 (Invitrogen, A-21036), 1:100 of Alexa fluor 568 phalloidin (Invitrogen, A12380), 1:100 of goat anti-rabbit Alexa fluor 488 (Invitrogen, A11008) and 1:1000 of 4,6-diamidino-2-phenylindol dihydrochloride (DAPI, Molecular Probe, D9542) in 0.5% (v/v) PBS-BSA, and 20μL was placed on each coverslip for 1 hour under a humid dark chamber. Back in the original well a second time, coverslips receive three final 0.5 % (v/v) PBS-BSA washes. Finally, a quick rinse with pure H2O is done before the mounting. The Prolong gold antifade reagent (Invitrogen, P36930) was used to mount the coverslips on glass slides. The cells would stay at room temperature overnight under a dark chamber before microscopy, and stored afterward at 4°C.

#### Microscopy

The microscope used was the Leica Thunder, an inverted fluorescence microscope. This microscope uses the built-in Thunder software to perform computational erasure, reducing background noise during acquisition. Thunder automatically considers optical parameters and provides continuous noise-free images.

Each condition was observed at x63 objective in immersion, with a 2×2 binning. Each field is the result of a 10×10 mosaic tile, randomly chosen around a position from which the focus was made to assure unbiased acquisition. Each tile was made with ten frames, with a scaling of 0,5μm between each frame. Four wave lengths were used from the higher to the lower:

- Vimentin: 635nm 400ms exposure, 30% laser power
- Actin: 560nm (filter over 590nm) 300ms exposure, 30% laser power
- ZO-1: 475nm, 400ms exposure, 30% laser power
- Nucleus: 390nm (filter over 440nm) 200ms exposure, 15% laser power

#### Segmentation

Maximum-Projection has been made on each image stack. Cell and nuclear contour segmentations were performed with cellpose using *cyto2*.^[23]^ for the cell labelling or *nuclei* for the nuclear labelling using the Napari user interface. The parameters used during each segmentation were:

Typical model:

- Diameter: Obtained digitally via image
- Cell Probability Threshold: -1.65
- Model Likeness Threshold: 23.84
- Average 4 nets: Checked
- Resampling Dynamics: Checked

Corrections were then made manually, if necessary, especially for the cell contours. Cells that overlap, or truncated at the edge of the field, have been removed.

#### Single-cell analysis

A macro has been designed for the single cell analysis on the open-source software FIJI.^[24]^ A Principal Component Analysis (PCA) is realized on segmented cell and nuclear contours by using celltool.^[25]^ For the UMAP analysis, we made the PCA of every segmented cell (and nucleus in a different analysis) at once. For the Celltool morphology map, we made the PCA with the three replicates, one cell line and one condition at a time. The previous two data sets were then exported to MS Excel and manually merged into one. The figures were made from these files with Python (v3.8.10) using the open-source user software Spyder.

#### Statistical analysis

Welch’s t-test analysis was performed due to the large sample size and different variance between conditions, to calculate P-value for all the experiments using Python. All four cell lines have been made in triplicate (N=3). *, **, ***, and **** represent P % 0.05, P% 0.01, P% 0.005, and P% 0.001 respectively. The bar graph is written by mean with Standard Deviation.

## Author contributions

Conceptualization, A.L F.C S.K; methodology, A.B.; investigation, A.B, A.L, and F.C; writing – original draft, A.B writing – review & editing, A.L F.C S.K. M.M, and AB; funding acquisition, A.L F.C; supervision, A.L, F.C; project administration, A.L, F.C.

## Acknowledgment

We would like to thank Dr. Juliette Griffié for her help in the statistical analysis suggestions. We also thank the whole MEC-uP group at ERRMECe Laboratory for all their feedback and help during the writing of this reports. This work is part of the Alphonse Boché doctorate thesis.

## Data and materials availability

All data needed to evaluate the conclusions in the paper are present in the paper and/or the Supplementary Materials. The original code generated during this study is available at [Interactive pipeline]: https://github.com/AlBoche/Image-based-single-cell-phenomics/tree/main. Cell acquisitions, segmentation contours and raw data are available at upon request, due to file size limitations.

