## Supplementary Materials for "Phenomics demonstrates cytokines additive induction of epithelial to mesenchymal transition"

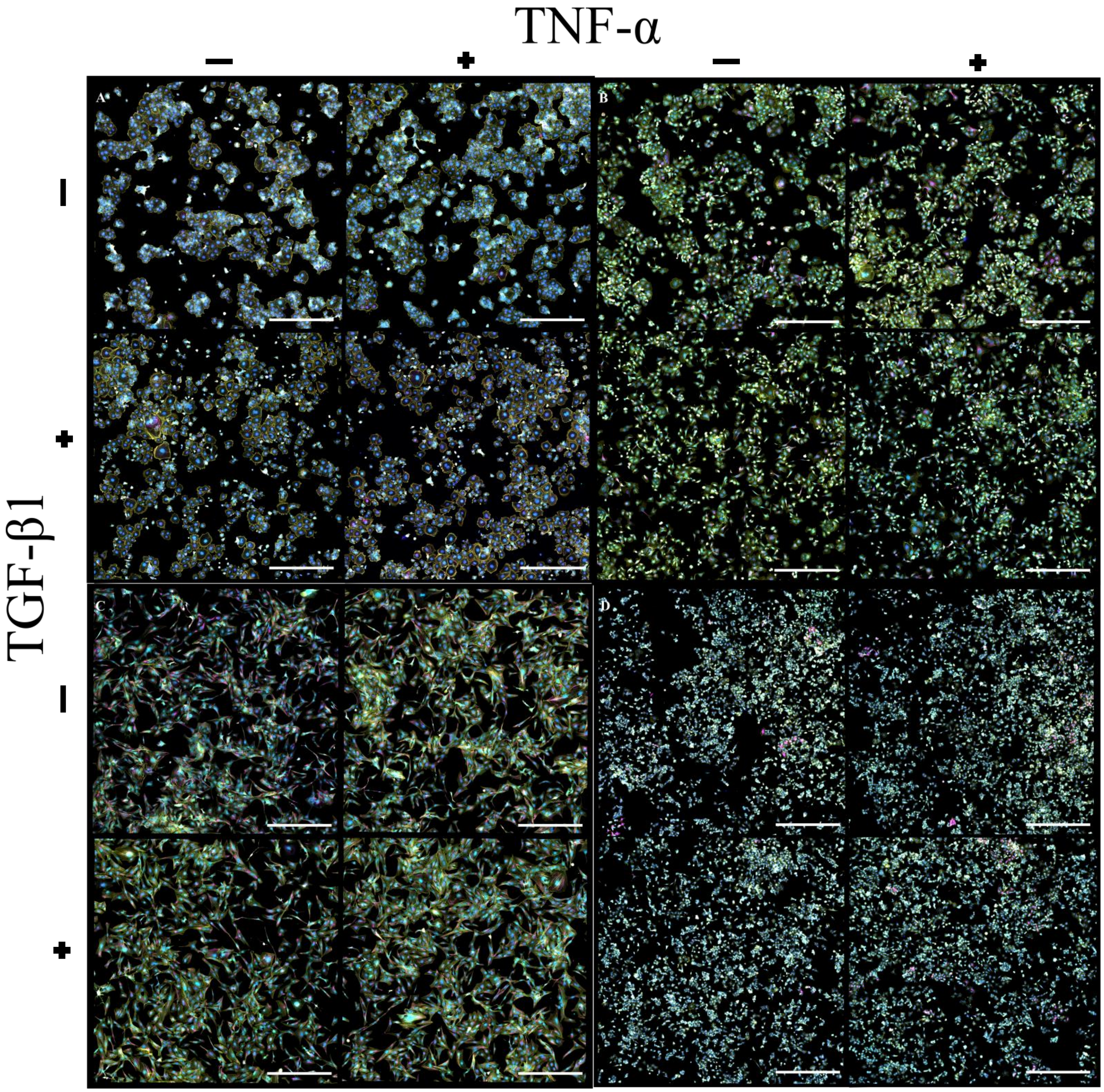

Figure S1: Full wide-scale replicate for microscopy experiment. A) OVCAR3, B) IGROV1, C) SKOV3, D) A2780_C10 are represented under four conditions: Control, TNF-α, TGF-β1 and TNF-α + TGF-β1 combination. The labels are blue for nucleus, cyan for ZO-1, yellow for actin and magenta for Vimentin. Scale bar = 500μm.

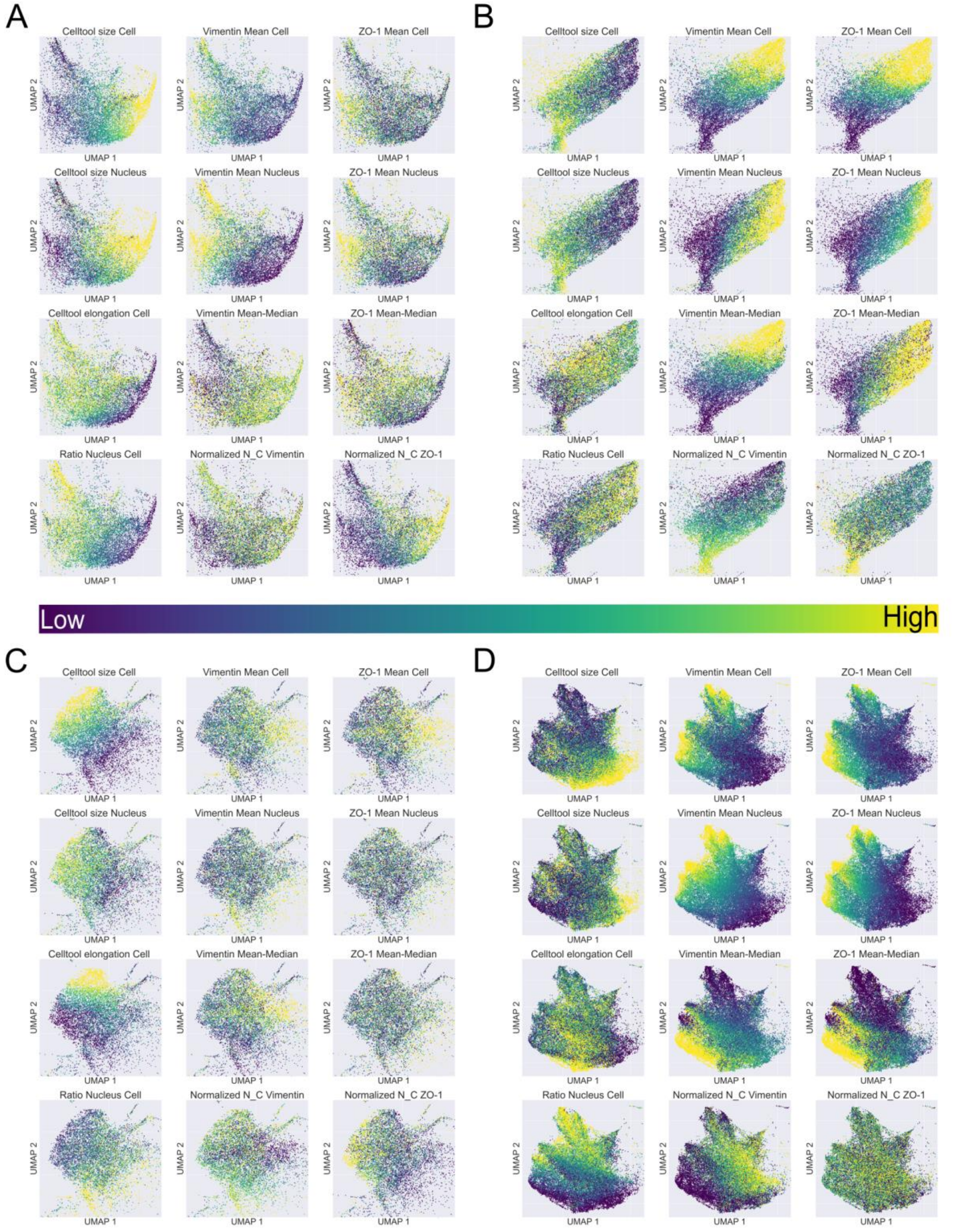

Figure S2 Heatmaps of single parameters after clustering. Four features for morphological, Vimentin and ZO-1 are represented. (A) OVCAR3 (B) IGROV1 (C) SKOV3 and (D) A2780_C10 are represented with the same classification.

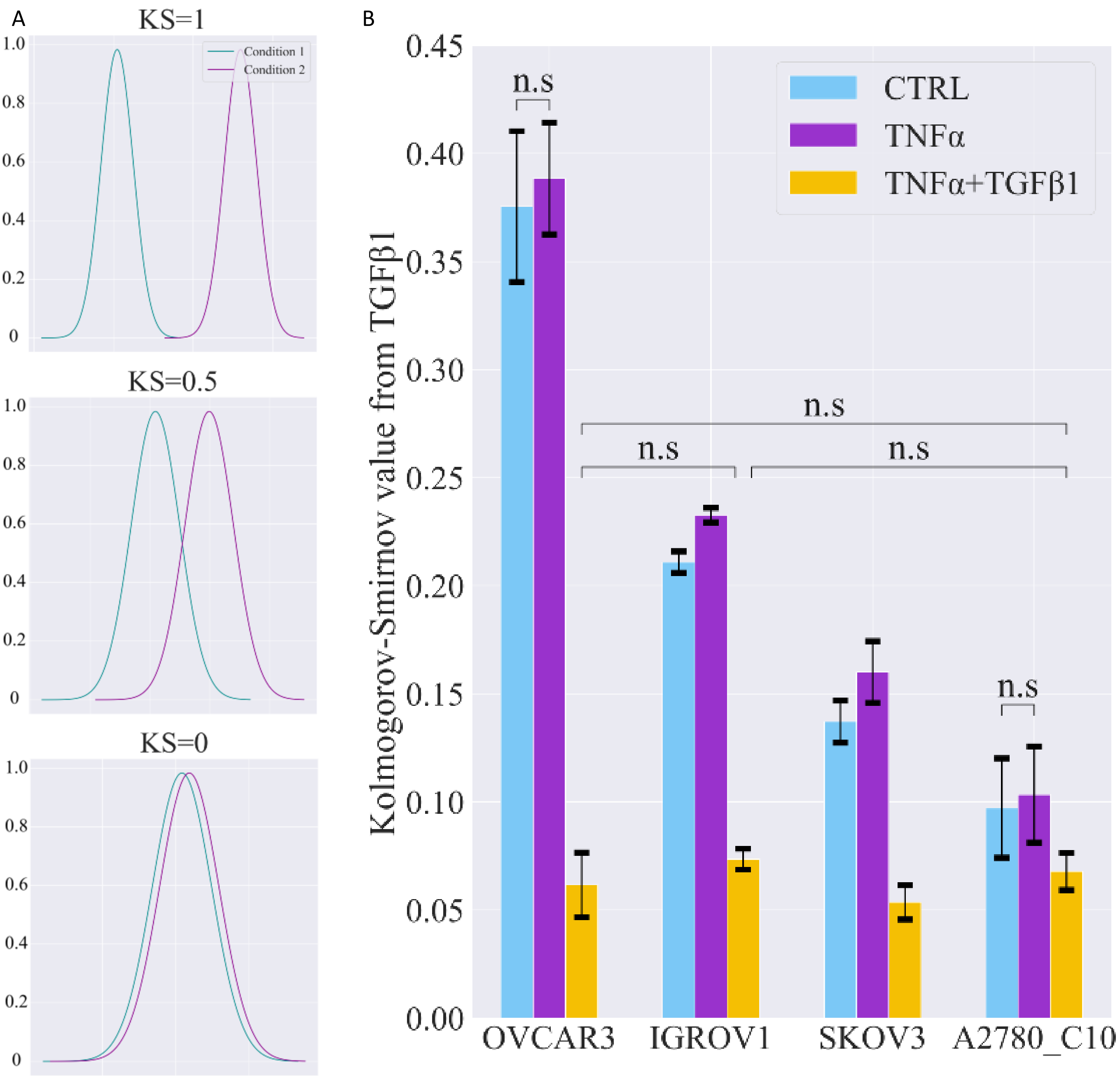

Figure S3: (A) Kolmogorov Smirnov to represent the difference between two distributions. A KS value of 0 represents two indissociable distribution, while a KS value of 1 represents two distinct distributions. (B) KS value from all conditions compared to the TGFβ1 condition. KS values are obtained after 50 generations of UMAP on all replicates. N=3. If not represented, significance is at least $p<0.05$.

| Group | Parameters | Description |
|---|---|---|
| Morphology | Area Cell | Area of cellular segmentation. |
| | Feret Cell | The longest distance within a segmented cell. |
| | Solidity Cell | Ratio of the Area with its convex hull. |
| | Area Nucleus | |
| | Perimeter Cell | Perimeter of the cellular segmentation. |
| | Perimeter Nucleus | Perimeter of the nucleus segmentation. |
| | Minimum Feret Cell | The shortest distance for a contour to go through. Also named the calliper diameter. |
| | Aspect Ratio Cell | Ratio of the major and minor axis. High AR means a more elongated/thin morphology. |
| | Aspect Ratio Nucleus | |
| | Solidity Nucleus | |
| | Ratio Nucleus Cell | |
| | Celltool size Cell | Celltool distribution for the size. A higher value means a larger Cell. |
| | Celltool elongation Cell | Celltool distribution for the elongation. A higher value means a longer Cell. |
| | Celltool size Nucleus | Celltool distribution for the size. A higher value means a larger Nucleus. |
| | Celltool elongation Nucleus | Celltool distribution for the elongation. A higher value means a longer Nucleus. |
| Vimentin / ZO1 | Mean intensity Cell | Mean of intensity for Vimentin or ZO1. Ratio of total fluorescence to the Area. |
| | StDev Cell | Standard deviation of pixels intensities generating the mean value. |
| | Mode Cell | Most frequent data appearing in the intensity distribution. |
| | Minimum intensity Cell | The minimum pixel intensity detected in the segmentation. |
| | Maximum intensity Cell | The maximum pixel intensity detected in the segmentation. |
| | Median intensity Cel | The median value of intensity. Half of values are above, and half are below. |
| | Skewness Cell | Third moment of order. Positive value means an asymmetric distribution to the right, negative to the left. |
| | Kurtosis Cell | Fourth moment of order. Positive value means a more peaked distribution, negative value a flatter. |
| | Mean intensity Nucleus | |
| | StDev Nucleus | |
| | Mode Nucleus | |
| | Minimum intensity Nucleus | |
| | Maximum intensity Nucleus | |
| | Median intensity Nucleus | The median value of intensity. Half of values are above, and half are below. |
| | Skewness Nucleus | |
| | Kurtosis Nucleus | |
| | Mean-median difference | Difference between the Mean and the Median. Presence of outliers increases this difference. |
| | Normalization from Nucleus to Cell | Linear regression coefficient of the mean intensity from the nucleus to the cell membrane. A lower value means a more important distribution around the nucleus. |

Table S1 : List and explanation of the data presented in Figure 2B.

| **OVCAR3** | CTRL | TNFα | TGFβ1 | TNFα+TGFβ1 |
|---|---|---|---|---|
| Percentage of cell surface in fields | 0,39 ± 0,02 | 0,42 ± 0,05 | 0,4 ± 0,06 | 0,47 ± 0,03 |
| Number of cells | 1047,67 ± 93,25 | 1172,33 ± 127,03 | 790,33 ± 32,19 | 810,67 ± 79,21 |
| Uncorrected cells | 370,33 ± 13,65 | 437,33 ± 54,78 | 476 ± 41,22 | 485,67 ± 38,59 |
| Corrected cells | 677,33 ± 88,44 | 735 ± 130,41 | 314,33 ± 10,69 | 325 ± 40,63 |
| Less than 3 corrections (over 10) | 636,67 ± 84,1 | 691,33 ± 113,39 | 301,67 ± 15,14 | 315 ± 37,99 |
| More than 3 corrections (over 10) | 40,67 ± 5,03 | 43,67 ± 17,9 | 12,67 ± 7,37 | 10 ± 2,65 |
| **IGROV1** | CTRL | TNFα | TGFβ1 | TNFα+TGFβ1 |
| Percentage of cell surface in fields | 0,38 ± 0,02 | 0,41 ± 0,03 | 0,38 ± 0,04 | 0,37 ± 0,03 |
| Number of cells | 1273 ± 54,25 | 1331,67 ± 59,81 | 1329 ± 211,21 | 1292,33 ± 150,31 |
| Uncorrected cells | 550 ± 41,68 | 554,67 ± 94,08 | 463,33 ± 60,14 | 468,67 ± 58,07 |
| Corrected cells | 723 ± 26,66 | 777 ± 38,2 | 865,67 ± 151,23 | 823,67 ± 95,2 |
| Less than 3 corrections (over 10) | 648,67 ± 28,04 | 679,67 ± 25,58 | 719 ± 119,88 | 688 ± 67,62 |
| More than 3 corrections (over 10) | 74,33 ± 4,93 | 97,33 ± 16,5 | 146,67 ± 37,23 | 135,67 ± 29,02 |
| **SKOV3** | CTRL | TNFα | TGFβ1 | TNFα+TGFβ1 |
| Percentage of cell surface in fields | 0,54 ± 0,05 | 0,55 ± 0,07 | 0,52 ± 0,03 | 0,58 ± 0,05 |
| Number of cells | 911 ± 139,31 | 957,67 ± 119,46 | 745,67 ± 29,67 | 878 ± 99,2 |
| Uncorrected cells | 194,67 ± 41,19 | 242,67 ± 38,84 | 139,67 ± 47,48 | 193,67 ± 53,98 |
| Corrected cells | 716,33 ± 105,83 | 715 ± 113,17 | 606 ± 19,08 | 684,33 ± 67,66 |
| Less than 3 corrections (over 10) | 551 ± 74,22 | 577,67 ± 84,91 | 500,67 ± 11,06 | 574 ± 36,59 |
| More than 3 corrections (over 10) | 165,33 ± 35,91 | 137,33 ± 33,29 | 105,33 ± 22,55 | 110,33 ± 36,86 |
| **A2780_C10** | CTRL | TNFα | TGFβ1 | TNFα+TGFβ1 |
| Percentage of cell surface in fields | 0,3 ± 0,03 | 0,33 ± 0,01 | 0,33 ± 0,04 | 0,34 ± 0,02 |
| Number of cells | 2553,67 ± 146,21 | 2802,67 ± 256,92 | 2738 ± 58,89 | 2899,33 ± 126,14 |
| Uncorrected cells | 1401,67 ± 166,58 | 1678 ± 273,66 | 1599,67 ± 182,44 | 1653,33 ± 179,58 |
| Corrected cells | 1152 ± 299,05 | 1124,67 ± 16,74 | 1138,33 ± 123,55 | 1246 ± 54,67 |
| Less than 3 corrections (over 10) | 1044,67 ± 256,86 | 1045,67 ± 30,6 | 1069,67 ± 108,54 | 1165,67 ± 68,02 |
| More than 3 corrections (over 10) | 107,33 ± 45,49 | 79 ± 13,86 | 68,67 ± 15,01 | 80,33 ± 24,38 |

Table S2: Summary table of image acquisition and corrections needed on the topography normalization tools.
